# Impacts of development and adult sex on brain cell numbers in the Black Soldier Fly, Hermetia illucens L. (Diptera: Stratiomyidae)

**DOI:** 10.1101/2022.01.25.477588

**Authors:** Meghan Barrett, R. Keating Godfrey, Emily J. Sterner, Edward A. Waddell

**Author notes:** Corresponding author: Meghan Barrett Department of Biology, Drexel University, 3245 Chestnut St, Philadelphia PA 19104.

## Abstract

The Black Soldier Fly (*Hermetia illucens*, Diptera: Stratiomyidae) has been introduced across the globe, with numerous industry applications predicated on its tremendous growth during the larval stage. However, basic research on *H. illucens* biology (for example, studies of their central nervous system) are lacking. Despite their small brain volumes, insects are capable of complex behaviors; understanding how these behaviors are completed with such a small amount of neural tissue requires understanding processing power (e.g. number of cells) within the brain. Brain cell counts have been completed in only a few insect species (mostly Hymenoptera), and almost exclusively in adults. This limits the taxonomic breadth of comparative analyses, as well as any conclusions about how development and body size growth may impact brain cell populations. Here, we present the first images and cell counts of the *H. illucens* brain at four time points across development (early, mid, and late larval stages, and both male and female adults) using immunohistochemistry and isotropic fractionation. To assess sexual dimorphism in adults, we quantified the number of cells in the central brain vs. optic lobes of males and females separately. To assess if increases in body size during development might independently affect different regions of the CNS, we quantified the larval ventral nerve cord and central brain separately at all three stages. Together, these data provide the first description of the nervous system of a popular, farmed invertebrate and the first study of brain cell numbers using IF across developmental stages in any insect.

## Introduction

The Black Soldier Fly (*Hermetia illucens*, Diptera: Stratiomyidae) is a tropical species native to Central and South America that has been introduced across the globe due to its industry applications (Marshall et al. 2015, Kaya et al. 2021). *H. illucens* larvae are actively being explored for use as livestock feed; fishmeal replacements; biodiesel; human, animal, and food waste management; and even as a source of sustainable human protein (Sheppard et al. 1994, Banks 2014, Widjastuti et al. 2014, Cheng et al. 2017, Julita et al. 2018, Chia et al. 2019, Lee et al. 2021, Hopkins et al. 2021). An estimated 200 billion larvae are reared annually as of 2020 (Rowe 2020). The recent ascent of *H. illucens* to a place of economic significance means that basic research on their biology is lacking. Particularly, there have been no studies of their central nervous system, although these data could aid in understanding both their development and behaviors.

Despite their small sizes, insect brains are capable of supporting remarkably diverse and sophisticated behaviors: learning, long-term memory, processing multimodal sensory information, navigation, foraging in complex terrain, tool use, courtship, nestmate chemical/facial recognition, and more (Pierce 1986, Detrain et al. 1999, Dukas et al. 2006, van Zweden & d’Ettorre 2010, Sheehan & Tibbets 2011, Ritzman et al. 2012, Giurfa 2015, Buehlmann et al. 2020). Understanding how these behaviors can be completed with such a small amount of neural tissue requires understanding processing power (e.g. number of cells) within the brain. Brain cell counts have been completed for a remarkably small number of insects due to methodological challenges recently resolved through the application of the isotropic fractionation (IF; Herculano-Houzel & Lent 2005) method to insect brains (Godfrey et al. 2021).

Development, diet, body size, cognitive abilities, metabolic limitations, and more have currently unexplored effects on brain size, as measured by cell number or density. *H. illucens* is a particularly excellent system for understanding how larval body size scales with brain cell development; larvae have incredible bioconversion rates (Surendra et al. 2016), with a greater than 50-fold change in body weight between the third and sixth larval instar (when fed standard *Drosophila melanogaster* food diets; Kim et al. 2010). In addition, while adult males and females do not appear obviously dimorphic in their external anatomy outside of females’ larger body sizes, behavioral variation related to mating (Tingle et al. 1975, Copello 1926, Julita et al. 2020) could generate sexual dimorphism in functionally discrete regions of the brain.

Here, we present the first images of the larval and adult *H. illucens* brain. We quantify changes in total brain cell number at four time points across development (early, mid, and late larval stages, and both male and female adults). We determine the number of cells in the central brain versus optic lobes of male and female adults (to look for sexual dimorphism in the visual system), and quantified the larval ventral nerve cord separately from the brain. Together, these data provide the first description of the nervous system of a popular, farmed invertebrate and close relation of *D. melanogaster*, and the first study of brain cell numbers using IF across developmental stages in any insect.

## Materials and Methods

### Rearing and Collection

For larval stages 1 and 4, black soldier fly eggs were obtained from ReptiWorms (Chico, California). 0-24 hour old larvae were placed on a standard *Drosophila* larval food recipe (as in Kim et al. 2010) in an incubator at 25 °C and 50% RH with 24 hours of darkness. Larvae were collected and stored whole in Prefer fixative at 0 - 24 hours (L1) and 10-11 days old (approximately L4).

L6 larvae were either reared and collected from the ReptiWorms population at 36-37 days old (L6), or obtained from Sympton Black Soldier Fly (College Station, Texas). L6 were immediately cut in half and stored in Prefer fixative. Additional larvae from the Sympton BSF population were kept at 26 - 29 °C, 35 - 60% RH, and 12 - 12 light-dark until eclosion. Adults were collected within 96 hours of eclosion and anesthetized in a jar with a cotton ball soaked in isoflurane, prior to the removal of the head capsule. Heads were stored in Prefer fixative for a minimum of three days before dissection.

L1 larvae were too small to be weighed accurately, but were assumed to weigh the same amount as an egg (25 μg; Dortsman et al. 2017). Wet masses of fixed L4 (n = 3) and L6 (n = 24) larvae were obtained to the nearest 0.01 mg on an analytical balance (Mettler Toledo AT261; Marshall Scientific, Hampton, NH, USA) prior to dissection; larvae were blotted dry of excess fixative before weighing. A subset of larvae from various instars were weighed before and after fixation, and fixation was found not to majorly impact wet mass (linear regression, [fixed mass] = 0.996 [pre-fixation mass], n = 27, R^2^ = 0.999). L1 and L4 larvae were also too small to have their head widths accurately recorded. The head width of L6 larvae was measured to the nearest 0.01 mm using digital calipers. Adult flies were sexed, weighed to the nearest 0.01 mg, and their head width at the widest point, and head height, were measured to the nearest 0.01 mm using digital calipers.

### Dissection & Isotropic Fractionation

Brains were dissected in Phosphate Buffer Solution (PBS; MP Biomedicals LLC). Adult optic lobes (OL) were separated from the central brain (CB), and the retinas were removed. Ultra Fine Clipper Scissors II (Fine Science Tools) were used to separate the larval ventral nerve cord (VNC) from the brain. Dissected brains were stored at 4 °C in PBS for 24 hours. Adult brains were carefully blotted dry with a Kimwipe and the OL and CB weighed separately to the nearest 0.01 mg on an analytical balance in 80% glycerol to minimize evaporation.

As in Godfrey et al. (2021), brain tissue was homogenized in a glass tissue homogenizer with sodium citrate and Triton X detergent solution, diluted with PBS. Nuclei were labelled with the fluorescent probe, SYTOX Green (ThermoFischer Scientific), then counted with a haemocytometer under epifluorescence using a 40x/0.65 M27 objective on a Zeiss Axioplan microscope. Twelve subsamples of each homogenized brain region were counted and averaged to provide a mean number of brain cells in that brain region. Cell density was obtained by dividing the mean number of nuclei for a given brain region by the mass of that brain region (assuming one nucleus per brain cell). Each adult’s OL and CB were counted separately (n = 12 females, 11 males). For L6 larvae, the VNC or brain for three individuals were homogenized simultaneously (due to the small cell numbers in each individual), and the final count divided by three (n = 7 samples of 3 individuals). L6 larvae for IF were pulled from both the Sympton and Reptiworms populations, which did not differ in mean body mass or final brain/VNC counts.

### Immunohistochemistry

Immunohistochemistry was used to obtain cell numbers for L1 and L4 larvae, and to obtain representative images of the three larval instars and adult brains. Brains were dissected as in the preceding section except the optic lobes and retinas were left attached to the central brain for adults and larval ventral nerve cords were left attached to the central brain. For larval samples the nervous system was rinsed in PBS then incubated in a small volume of SYTOX Green (1:5000 in PBS) for 30 minutes. The nervous system was then cleared in increasing steps of glycerol (40%, 60%, 80% in PBS) at 35 °C and mounted on a slide with a polyvinyl alcohol mounting medium, Mowiol® 4-88 (Sigma-Aldrich), and covered with a #1.5 coverslip. Adult brains were embedded in 10% low-melting agarose and sectioned at 100 μm on a vibratome. Sections were then processed and mounted as described for larval brains. Samples were imaged on a Zeiss LSM 880 inverted confocal microscope. Larval brains were imaged using a 40x/1.3 Oil DIC M27 objective with optical section thickness of 1 μm (L1), 2 μm, (L4) or 3 μm (L6). Adult brains were imaged using a 20x/0.8 M27 lens with images acquired at an optical section thickness of 8 μm. Two observers counted each L1 and L4 (n = 2/instar) brain (VNC separately from the brain) and their counts were averaged.

### Statistical Analysis

GraphPad Prism v. 9.1.2 (GraphPad Prism for Windows 2021) was used for all statistical analyses. Shapiro-Wilk normality tests and an F-test for equal variance were used to determine if data met the assumptions of parametric tests. An unpaired t-test was used to analyze categorical differences between adult males and females in body mass, head width, head height, relative OL mass, and total brain cell numbers. Linear regressions were used to analyze the relationship between body mass and head width/height, brain mass (of males and females), OL mass (of males and females), CB mass (of all adults), and CB, OL, and total brain cell densities (of all adults). A quadratic regression was used to analyze the relationship between body and relative brain mass, and the fit was compared to a linear regression. Due to unequal variance, a Welch’s ANOVA with Dunnett’s T3 multiple comparisons test was used to assess differences in the number of cells in male and female OLs and CB; a one-way ANOVA with Bonferroni MCT was used to assess differences in the cell densities of male and female OLs and CBs.

## Results

### Description of the Larval Brain

The larval brain consists of two attached, spherical lobes, similar to *Drosophila melanogaster* (Figure 1). The lobes of the brain each connect to the first of the twelve segmented ganglia of the VNC. The twelve ganglia of the VNC each innervate one of the twelve body segments (including the head) of the larvae; each ganglion is composed of two, symmetrical regions that presumably innervate the left and the right sides of the body. While the VNC extends through nearly the entire body of an L1, it only extends through body segments 3 – 6 of the much larger L6.

**Figure 1.**
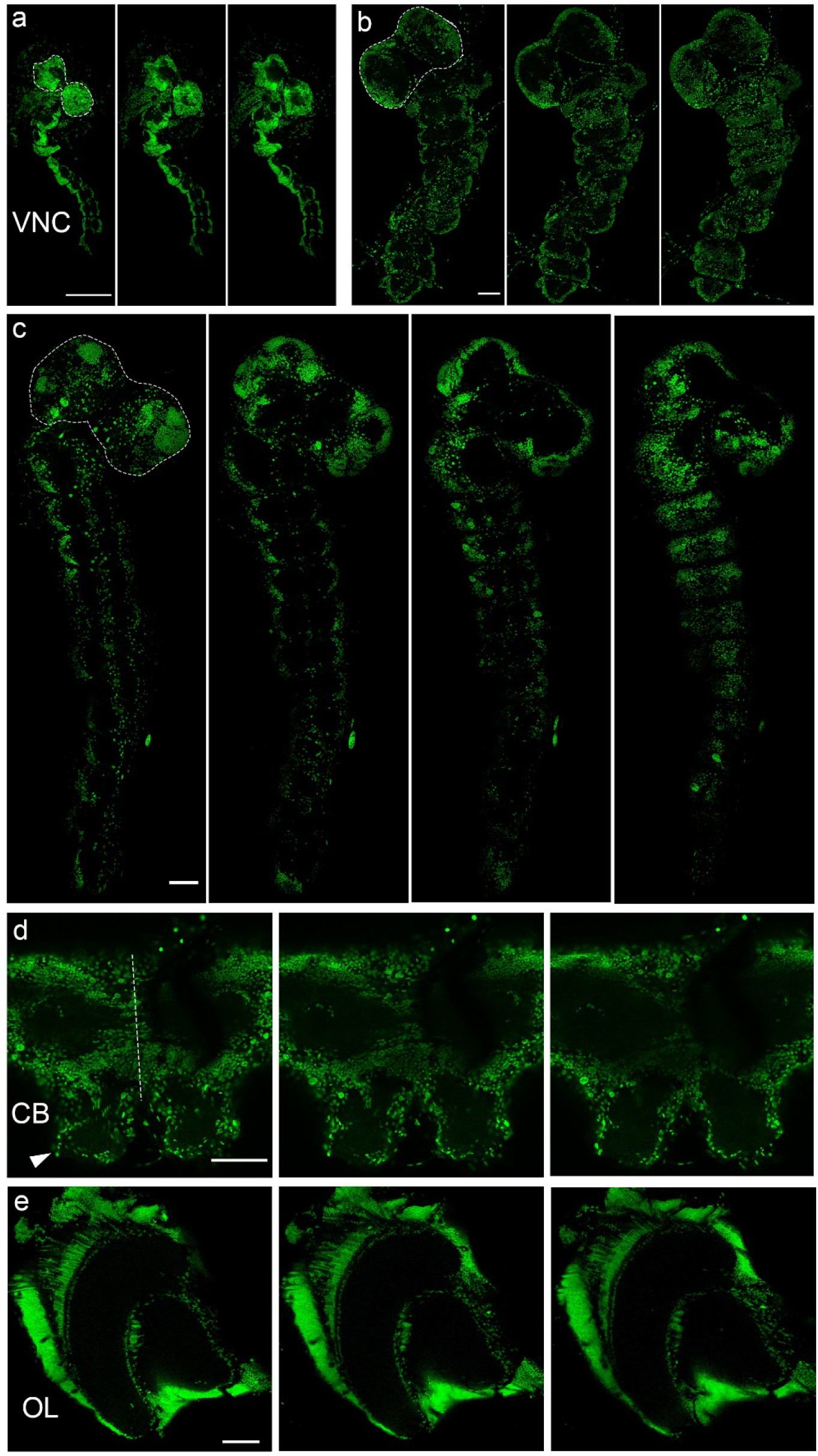
Central nervous system development in the Black Soldier Fly. Subsections of SYTOX Green labelled tissue imaged at 40x from (a) first (L1), (b) fourth (L4), and (c) sixth (L6) instar nervous systems showing the brain (dotted outline) and ventral nerve cord (VNC). Subsections from the adult (d) central brain (CB) and (e) optic lobes (OL) imaged at 20x and labelled with SYTOX Green. In (d) dotted line denotes brain midline and arrow indicates left antennal lobe. Scale bars = 100 μm.

### Changes in Total Brain Cell Numbers Across Development

There was an 8.8-fold increase in total brain cell number (from 2,314 ± 71 to 20,355 ± 2,780) with a 257-fold change in body mass across larval development (Table 1). During pupation, as the brain developed functionally discrete regions (Figure 1), flies produced a 16.2-fold increase in brain cell numbers (330,737 ± 62,376; Table 1).

**Table 1.** Changes in brain cell number across developmental stages and body mass.

| Life stage (mean body mass, mg) | Mean total number of brain cells ± SD | Method (n) |
| --- | --- | --- |
| L1 (0.025) | $2,314 \pm 71$ | IHC + Counting (n = 2) |
| L4 ($15.32 \pm 4.02$) | $11,529 \pm 1,294$ | IHC + Counting (n = 2) |
| L6 ($64.18 \pm 9.03$) | $20,355 \pm 2,780$ | IF (n = 7 samples, 3 individuals/sample) |
| Adult ($37.98 \pm 8.15$) | $330,737 \pm 62,376$ | IF (n = 23) |
| F only ($41.23 \pm 8.52$) | $299,011 \pm 59,519$ | IF (n = 12) |
| M only ($34.44 \pm 6.32$) | $365,347 \pm 46,233$ | IF (n = 11) |

The number of cells in the larval VNC increased more slowly brain mass across developmental stages (Figure 2A). L1 have 2,684 ± 186 cells, L4 have 12,249 ± 1,153, and L6 have 16,146 ± 1,386 cells in the VNC, a 6-fold increase in VNC cell number across development.

**Figure 2.**
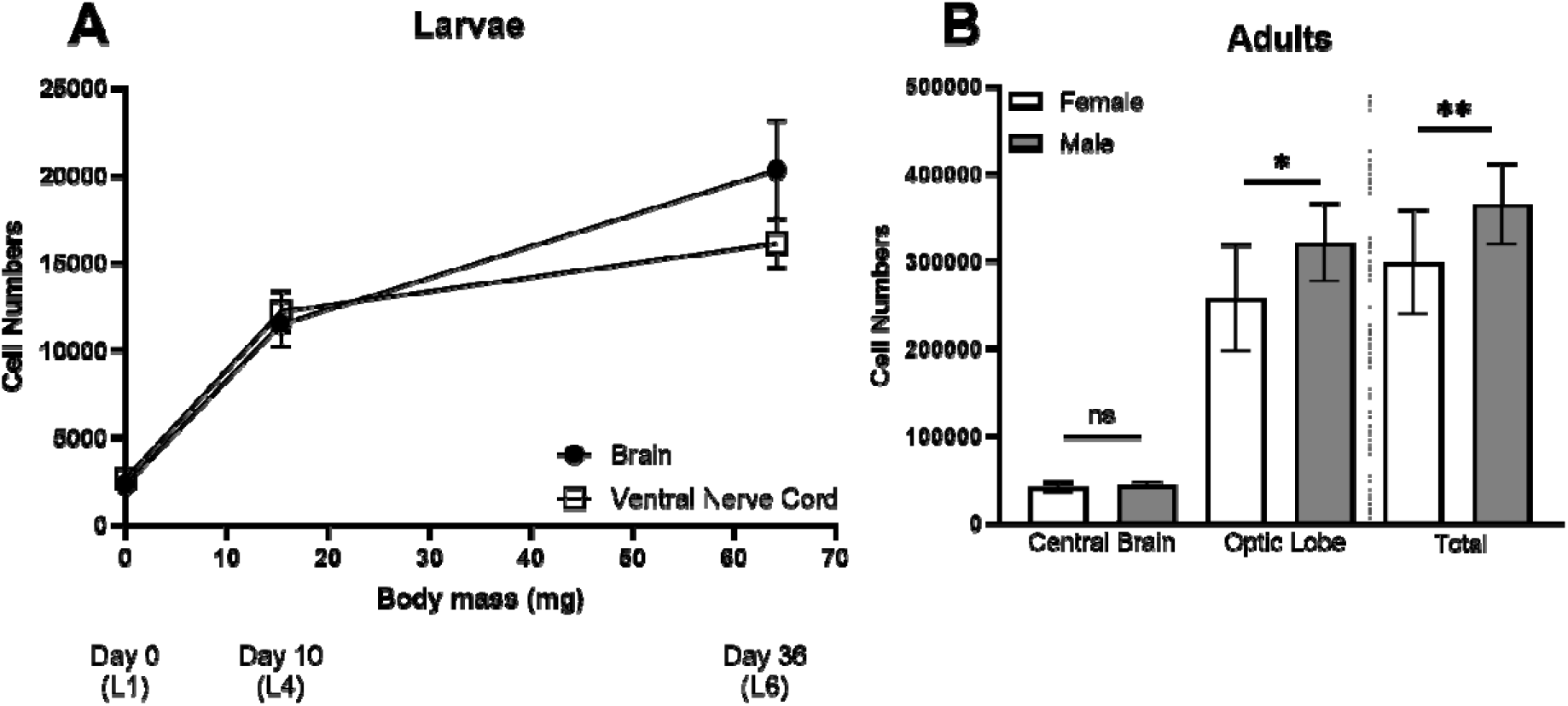
Brain cell numbers during larval development and in male vs. female adult Black soldier flies. A) Black soldier fly larvae have increased brain and ventral nerve cord cell numbers. Brain and ventral nerve cord cell numbers increase faster, compared to body mass, earlier in development (L1 – L4) as opposed to later in development (L4 – L6). B) Adult, male *H. illucens* have an increased total number of brain cells compared to females (Unpaired t-test: t = 2.97, df = 21, p = 0.0074). This difference is driven by differences in the optic lobes (Welch’s ANOVA: W = 180.20, df = 20.30, p < 0.0001; Dunnett’s T3 MCT, OL: t = 2.91, df = 20.12, p = 0.0171), as males and females have the same number of cells in the central brain (CB: t = 1.00, df = 17.74, p = 0.55). ns = not significant; * = p < 0.05; ** = p < 0.01

### Body Size Effects and Sexual Dimorphism in Adults

Males in our sample had reduced mean body sizes compared to females, as measured by mass (Unpaired t-test, t = 2.15, df = 21, p = 0.043; males, range: 22.15 – 42.42 mg; females, range: 32.30 – 56.43 mg), but not head width (t = 1.69, df = 21, p = 0.10) or head height (t = 0.78, df = 21, p = 0.45). Head width and head height scaled hypoallometrically with adult body mass, suggesting that larger head sizes are prioritized at small body sizes (HW: log [head width] = 0.45 log [body mass^1/3^] + 0.28, F =32.83, df = 21, p < 0.0001, R^2^ = 0.61; HH: log [head height] = 0.31 log [body mass^1/3^] + 0.14, F =7.97, df = 21, p = 0.01, R^2^ = 0.27).

Across adults, relative brain mass increase non-linearly with decreasing body mass in accordance with Haller’s rule ([Brain:body mass] = 1.23 e^-05^ [body mass]^2^ – 0.001 [body mass] + 0.04, F = 7.20, df = 20, p = 0.0143, R^2^ = 0.61). Female and male brain mass scaled hypoallometrically with body mass, but at different rates (F = 7.43, df = 19, p = 0.0134); female brains increased in mass as body size increased, while male brain mass was unaffected by increasing body mass (females: log [brain mass] = 0.51 log [body mass] – 1.07, F = 7.33, df = 10, p = 0.022, R^2^ = 0.42; males: F = 0.91, df = 9, p = 0.37).

The relationship between brain and body mass was driven by differences in OL scaling, as adult OLs accounted for 74% ± 4% of total brain mass (no difference in relative OL mass in males and females; Unpaired t-test: t = 0.19, df = 21, p = 0.85). Female OLs scaled hypoallometrically with body mass, while male OL mass was unaffected by body size (females: log [OL mass] = 0.52 log [body mass] – 1.20, F = 13.70, df = 10, p = 0.0041, R^2^ = 0.58; males: F = 0.58, df = 9, p = 0.47). CB mass was unaffected by body size in adults (F = 2.36, df = 21, p = 0.14).

Males had significantly more brain cells than females (Table 1, Figure 2B; Unpaired t-test: t = 2.97, df = 21, p = 0.0074). This difference was due to increased numbers of cells in their optic lobes, but not the central brain region (Figure 2B; Welch’s ANOVA: W = 180.20, df = 20.30, p < 0.0001; Dunnett’s T3 MCT, OL: t = 2.91, df = 20.12, p = 0.0171; CB: t = 1.00, df = 17.74, p = 0.55). Males had 321,776 ± 44,636 cells in their optic lobes compared to 257,566 ± 60,579 for females. Adults had 42,462 ± 5,222 cells in the central brain region.

Across adults, cell density (nuclei/mg) in the OL and total brain decreased as body mass increased, while CB density remained statistically similar, but trended towards decreasing (Total: [brain cell density] = -9405 [body mass] + 996185, F = 6.23, df = 21, p = 0.021, R^2^ = 0.23; OL: [OL cell density] = -8300 [body mass] + 872429, F = 5.27, df = 21, p = 0.0322, R^2^ = 0.20; CB: F = 4.19, df = 21, p = 0.054). Males had more dense brains than females in the OL but not the CB (ANOVA: F = 40.41, df = 42, p < 0.0001; Bonferroni MCT, OL: t = 6.10, df = 42, p < 0.0001; CB: t = 1.79, df = 42, p = 0.16).

## Discussion

In this study, we provide the first images of the *H. illucens* brain across development. In addition, we determine how developmental stage impacts brain cell numbers, both across larval stages and following metamorphosis, completing the first intraspecific developmental comparison of brain cell numbers using IF. Finally, we separately compare the central brain and optic lobes of male and female *H. illucens*, and find sexual dimorphism in the adult OLs but not the CB, which has not been reported in other Diptera.

First instar *D. melanogaster* larvae have 2,000 cells in the brain – roughly comparable to *H. illucens* (Scott et al. 2001, Nassif et al. 2002, Avalos et al. 2019). Third (final) instar *D. melanogaster* larvae have an estimated 8-10,000 cells in the brain (Nassif et al. 2002, Thum & Gerber 2019); this appears roughly comparable to the third stage of *H. illucens* (as the fourth stage brain contained ∼13,000 cells), suggesting there may be certain common developmental rules may regulate neuronal differentiation during larval molts in this region across some species. *H. illucens* have another three molts before pupation, and the brain reaches 20,000 cells by the final L6 stage.

*H. illucens* adults have two to three times the number of protocerebral brain cells as *D. melanogaster* (most *D. melanogaster* counts range from 93,000 through 133,000; Godfrey et al. 2021, Scheffer et al. 2020, Mu et al. 2022; but some estimate 208,000 cells, Raji & Potter 2021). The vast majority of this increase in cell number is likely due to the optic lobes. *D. melanogaster* have around 25,000 cells in the CB (Scheffer et al. 2020; though Mu et al. 2022 estimate 43,000, and Raji & Potter 2021 estimate 101,000), as compared to our estimate of 42,000 in *H. illucens* adults. OL cell number estimates in *D. melanogaster* (90,000 in Mu et al. 2022; 107,000 in Raji & Potter 2021) are much lower than the 250,000 and 320,000 cells in the OLs of female and male *H. illucens*, respectively. OL cell nuclei were noticeably smaller than those in the CB (Figure 1) similar to *D. melanogaster* (Mu et al. 2022).

Quality of the larval diet is known to impact neurogenesis – reduced diet quality decreases the number but not diversity of cells in adult visual centers in *D. melanogaster* (Lanet et al. 2013). Body size is linked to diet quality, rearing density, and temperature in *H. illucens* (Chia et al. 2018, Barragan-Fonseca et al. 2018, Jones & Tomberlin 2019, Gobbi et al. 2013, Addeo et al. 2021), and growth and development time can also vary significantly between populations of insects (Edgar 2006, Zhao et al. 2013). It is possible that diet, temperature, or source population may affect the total number of brain cells estimated for an insect species.

The increased number of OL cells in adult males is surprising. There is no obvious sexual dimorphism in eye morphology between males and females (as in honey bees, for example; Streinzer et al. 2013). Other Dipterans (such as *D. melanogaster*, or the mosquito species: *Aedes aegypti, Anopheles coluzzii*, and *Culex quinquefasciatus*; Raji & Potter 2021) do not demonstrate differences in optic lobe cell numbers between males and females. In natural conditions, *H. illucens* gather in leks to chase after and mate with females. Males often engage in aggressive, territorial, or courtship interactions with other males (Tomberlin & Sheppard 2001, Giunti et al. 2018). However, the sensory signals used by males to distinguish receptive females vs. unreceptive males are still unclear. Many male insects use a combination of chemosensory, acoustic, or visual cues to locate females (Benelli et al. 2014, Bonduriansky 2001). In *H. illucens*, acoustic signals are likely necessary for male courtship initiation (Giunti et al. 2018). However high-intensity light conditions with specific spectral characteristics have proven to be absolutely critical for encouraging mating in both captive and outdoor populations (Oonincx et al. 2016, Tomberlin & Sheppard 2002, Tingle 1975, Liu et al. 2020, Zhang et al. 2010, Macavei et al. 2020, Klüber et al. 2020, Heussler et al. 2018, Holmes 2010, Nakamura et al. 2016, Schneider 2020). This behavioral data, supported by the increased number of brain cells we found in the optic lobes of males, suggests that visual cues may also be very important for mediating some aspects of male mating behaviors.

This study is the first to describe the structure of the BSF larval and adult central nervous systems. Our results provide evidence for patterns of larval brain cell development in a second Dipteran species and demonstrate the use of IF for intraspecific comparisons across and within life stages. Our data suggest there is sexual dimorphism in the OLs of adults, which supports previous behavioral data demonstrating the importance of light conditions for BSF mating behaviors. Overall, our study suggests that IF can be used to more easily determine developmental patterns of brain complexity (as measured by brain cell number) in a wider variety of arthropod taxa, as well as intraspecific variation due to sexual dimorphism, age, diet, developmental conditions, and more.

## Acknowledgements

The Stanford Laboratory at Drexel University provided use of their incubator for specimen rearing. Wulfila Gronenberg provided extra reagents and equipment, as well as essential methodological guidance. We would like to thank Patty Jansma at the University of Arizona Imaging Core Optical facility for her microscopy advice and support.

